# Semi-supervised deep learning with graph neural network for cross-species regulatory sequence prediction

**DOI:** 10.1101/2022.05.17.492285

**Authors:** Raphaël Mourad

**Affiliations:** MIAT, INRAE, 31326 Castanet-Tolosan, France; University of Toulouse, UPS, 31062 Toulouse, France

**Keywords:** Regulatory genomics, Deep learning, Semi-supervised learning, Graph neural network

## Abstract

Genome-wide association studies have systematically identified thousands of single nucleotide polymorphisms (SNPs) associated with complex genetic diseases. However, the majority of those SNPs were found in non-coding genomic regions, preventing the understanding of the underlying causal mechanism. Predicting molecular processes based on the DNA sequence represents a promising approach to understand the role of those non-coding SNPs. Over the past years, deep learning was successfully applied to regulatory sequence prediction. Such method required DNA sequences associated with functional data for training. However, the human genome has a finite size which strongly limits the amount of DNA sequence with functional data available for training. Conversely, the amount of mammalian DNA sequences is exponentially increasing due to ongoing large sequencing projects, but without functional data in most cases. Here, we propose a semi-supervised learning approach based on graph neural network which allows to borrow information from homologous mammal sequences during training. Our approach can be plugged into any existing deep learning model and showed improvements in many different situations, including classification and regression, and for different types of functional data.

## 1 Introduction

Complex diseases are caused by the combination of multiple gene mutations with lifestyle and environmental factors [1]. Complex diseases are frequent in the population and include Alzheimer’s disease, autoimmune diseases, asthma and some cancers [2–4]. Over the past decade, genome-wide association studies (GWASs) have systematically identified thousands of single nucleotide polymorphisms (SNPs) associated with these diseases [1]. However, an important limitation of GWASs comes from the difficulty of gaining insight into the underlying biological mechanism, since over 95% of associated SNPs are located outside coding sequences [5]. Interestingly, the majority of the non-coding SNPs (75%) are located within regulatory elements [5], suggesting an important role for these SNPs in the deregulation of gene expression.

Deep neural networks have achieved state-of-the-art performance for the prediction of regulatory elements from DNA sequences. The first models were based on the stacking of one or few convolutional layer(s) followed by a dense network, thereby capturing combinations of DNA motifs specific to DNA binding proteins [6,7]. However, these initial models could not model long-range relations among DNA motifs within the sequence, and thus missed the complex grammar of the regulatory genome. To tackle this issue, several approaches were proposed such as recurrent neural networks with LSTM for few hundred bases [8], dilated convolutions [9] and transformers [10] for hundreds of kilobases. These models were trained using labeled data (supervised learning), but recently new models were trained without labeled data (unsupervised learning) [11,12].

Semi-supervised learning trains a model using not only labeled data generally available in small amount, but also using unlabeled data often available in large amount [13]. Graph neural networks (GNNs) were recently proposed for semi-supervised learning graph-structured data that is based on an efficient variant of convolutional neural networks which operate directly on graphs [14]. Instead of training individual embeddings for each node of the graph, a GNN aggregates features from a node’s local neighborhood, thereby borrowing information from unlabeled nodes. In the simplest GNN, the graph convolutional network (GCN), the aggregation simply consists in a linear combination from a node’s local neighborhood embedding [14]. Many improvements of the GCN were later proposed, including approximate personalized propagation of neural predictions (APPNP) with an improved propagation scheme based on personalized PageRank [15], GraphSAGE which concatenates the aggregation and the original node’s embeddings [16] and Graph Attention layer (GAT) [17] where edges have weights computed by an attention mechanism.

Here, we propose a novel framework for regulatory sequence prediction using semi-supervised learning with graph neural network. For this purpose, neural networks are trained not only from labeled sequences (*e.g*. human genome with ChIP-seq experiment), but also from unlabeled sequences (from other species without ChIP-seq experiment, *e.g.* chimpanzee, rabbit or dog). In order to incorporate unlabeled sequences in the training, a graph neural network connecting homologous sequences is used. Compared to supervised learning, the proposed semi-supervized learning allows to train models from a much larger number of sequences, without needing additional experiments since many unlabeled (unannotated) genomes are already available, while the vast majority of functional experiments such as ChIP-seq are only available in human, or to some extent in mouse.

## 2 Materials and Methods

### 2.1 Human experimental data

We used publicly available CTCF, ESR1, POL2, H3K4me3 ChIP-seq data and input data of human lymphoblastoid GM12878 from Gene Expression Omnibus (GEO) accession GSE31477 and GSE170139 from ENCODE [18]. We used publicly available ATAC-seq data of lymphoblastoid GM12878 from Gene Expression Omnibus (GEO) accession GSE170918 from ENCODE [18]. All the data was mapped on hg38 and peak calling was done using macs3 [19].

### 2.2 Mouse experimental data

We used publicly available CTCF, POL2 and H3K4me3 ChIP-seq data and input data of mouse lymphoblastoid CH12 (GM12878 analog) from Gene Expression Omnibus (GEO) accession GSE49847 from Mouse ENCODE [20]. All the data was mapped on mm10 and peak calling was done using macs3 [19].

### 2.3 Labeled DNA sequences

We binned the labeled genome (*e.g.* human genome assembly hg38) into non-overlapping genomic intervals of 200 b, where a given bin was considered as either bound by a TF (if overlapping > 50%), or not bound otherwise (classification setting). For each 200 b bin, we extracted the surrounding DNA sequence of 1 kb (containing the bin and the context).

### 2.4 Homologous DNA sequences

Given a labeled bin sequence (*e.g.* from hg38), the homologous sequences in other species’ genomes (*e.g.* from panTro6, oryCun2 and canFam3) were mapped using the following approach. We first mapped the labeled bin to another species genome using liftover. If a labeled bin mapped to different loci in another species genome but the loci were close to each other (separated by less than 50 b), the different loci were merged into one loci. Then the DNA sequence of 1 kb surrounding the loci was extracted and considered as a homologous sequence. If the bin mapped to different and distant loci (> 50 b) or if the bin did not map to any loci, then no homologous sequence was extracted (the homologous sequence was filled with N’s).

To search for homologous sequences, we used the following mammalian genomes: Rhesus monkey (rheMac10), marmoset (calJac3), chimpanzee (panTro6), pygmy chimpanzee (panPan2), Sumatran orangutan (ponAbe3), gorilla (gorGor6), olive baboon (papAnu4), crab-eating macaque (macFas5), Bolivian squirrel monkey (saiBol1), northern white-cheeked gibbon (nomLeu3), gray mouse lemur (micMur2), small-eared galago (otoGar3), mouse (mm10), rat (rn7), ferret (musFur1), rabbit (oryCun2), pork (susScr11), cat (felCat9), dog (canFam3), horse (equCab3), cow (bosTau9) and opossum (monDom5).

### 2.5 Model architecture

The proposed model is illustrated in Figure 1. The model takes two inputs: i) a set of 1 kb DNA sequences in one species (for instance, human) and the corresponding homologous sequences in other species (for instance, chimpanzee, mouse and dog), and ii) a set of graph matrices connecting the homologous sequences from the different species. As output, the model predicts either the probability of a DNA sequence to bind a given TF within a 200 b DNA sequence centered on the 1 kb DNA sequence.

**Figure 1.**
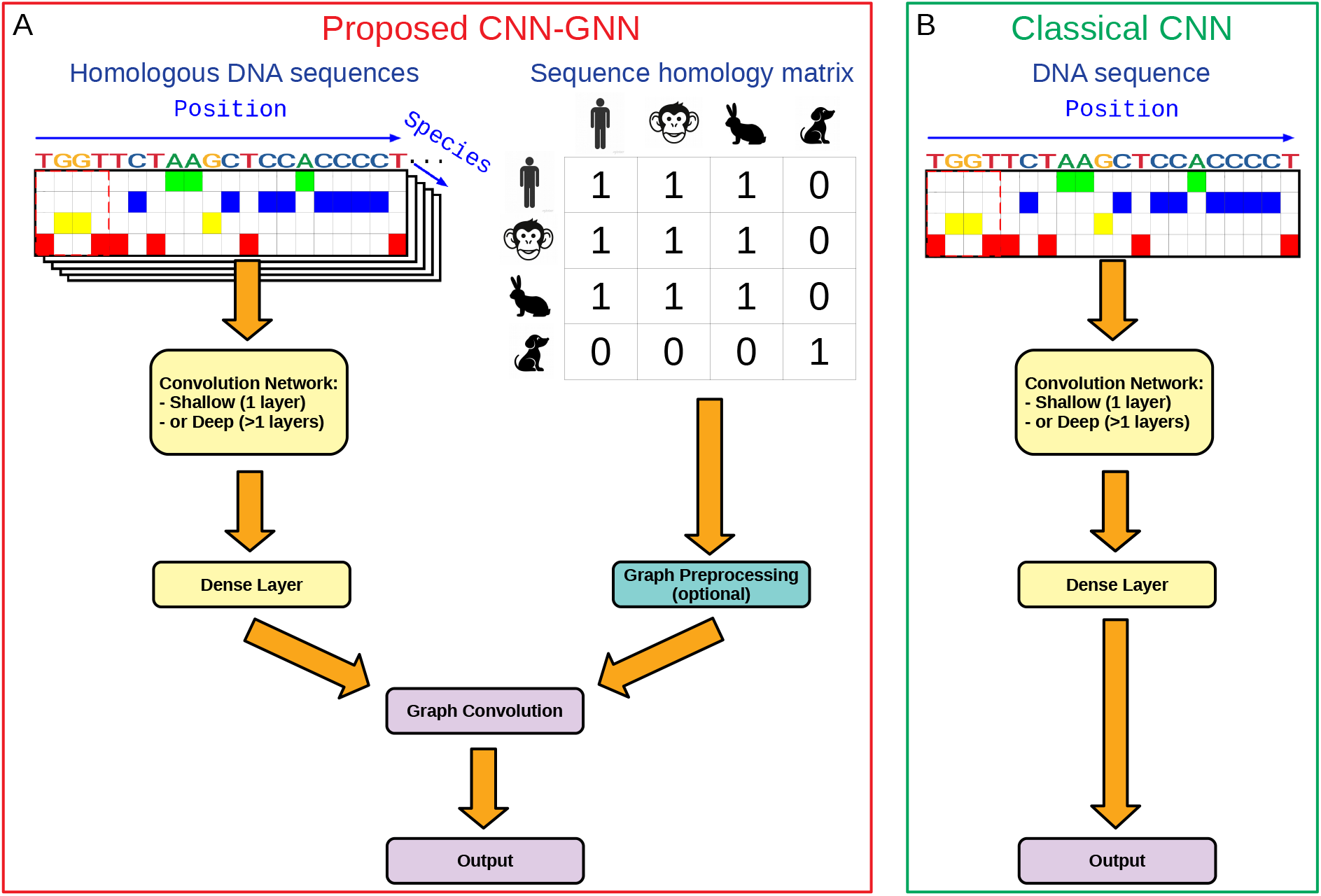
Sketch of the proposed semi-supervised model, implementing a convolutional network within a graph neural network (so called CNN-GNN), and comparison with the classical convolutional network (CNN).

As done in state-of-the-art, our model first models the DNA sequences using a convolutional network (left side of the network, Figure 1A). The convolutional neural network (CNN) can be composed of only one convolutional layer (shallow convolutional network). Alternatively, the convolutional network can be composed of a convolutional layer followed by dilated convolutions with residual connections (deep convolutional network), providing better performances in most situations by modeling long-range relations. Each convolutional layer comprises 64 filters (also called kernels). In the first layer, filters of size=24 were used. In the next layers (present in the deep convolutional network), filters of size=3 with an increasing dilation rate of 2 for each additional layer were used instead. The convolutional network is followed by a global maximum pooling and a dense layer of 10 neurons. If graph convolution was not used (classical CNN), then a last layer of 1 neuron was used to predict the outcome. Alternatively, graph convolution was used as described in the paragraph bellow.

In parallel, the model takes as a secondary input the graph matrix connecting homologous sequences between species (right side of the network, Figure 1A). Depending on the graph convolution used, the graph matrix can be preprocessed. For instance, for the GCN, APPNP or GSC convolutions, the graph matrix is preprocessed as follows:

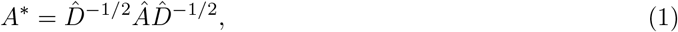

where 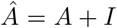 is the adjacency matrix with added self-loops (*I*) and 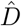 is its degree matrix. For the GraphSAGE or GAT convolutions, no graph matrix preprocessing is needed. Lastly, a graph convolution is used to convolve the activation values from the dense layer over the sequence homology matrix producing sequence output with 1 neuron. We chose the GraphSAGE convolution for our model as providing the best results, although we compared different graph convolutions in the Section Results and Discussion, Subsection Comparison with the baseline model, Supp Fig S1. For the GraphSAGE, graph convolution was done over the dense layer of 10 neurons as follows:

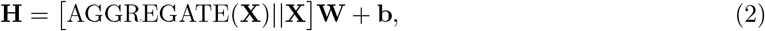

where AGGREGATE aggregates a node’s neighborhood by summing, *i.e.* it aggregates the features over the different homologous sequences. A last layer of 1 neuron then combines the aggregated features and predicts the outcome. Because the proposed model combined a GNN on the top of a CNN, we called it “CNN-GNN”. We compared the CNN-GNN with the baseline CNN where graph convolution was not implemented (Figure 1B).

### 2.6 Model training

Model training was done using the following hyper-parameters: ADAM optimizer, a minimum of 5 epochs, a maximum of 30 epochs, a patience of 3 epochs, a learning rate = 0.0005, a clipping norm = 0.0001, a batch size=100. For classification, binary cross-entropy was used as loss, where as for regression, the Poisson loss was used instead. The models were trained with 100 thousand sequences in balanced manner: 50 thousand bin sequences corresponding to observed ChIP-seq peaks (or other experimental peaks) and 50 thousand randomly drawn bin sequences. When the number of peaks was not enough to obtain 50 thousand bin sequences, upsampling was used to reach 50 thousands.

### 2.7 Prediction of SNP effect

We used the deep learning models to predict the effect of SNPs on functional data such as protein binding or chromatin accessibility. To compute the effect of a SNP, we used the following approach for classification as in [6]. First, model predictions were computed both for the DNA sequence comprising the reference SNP allele (*p_ref_*) and the same DNA sequence but with the alternative SNP allele (*p_alt_*). Then, the SNP effect was computed as (*p_alt_* – *p_ref_*) · *max*(0, *p_alt_, p_ref_*), where *p_alt_* is the prediction from the alternative allele and *p_ref_* the prediction from the reference allele.

To predict SNP effect with the CNN-GNN, graph convolution was not used for prediction (only used for training), since it is not possible to accurately know the alternative and reference alleles in homologous sequences from other species. Hence, the sequence homology matrix values were set up to zeroes, except for the labeled sequence and itself (first row and first column of the matrix). Then the SNP effect was computed as detailed in the previous paragraph.

### 2.8 Implementation and Availability

The models were developed using Tensorflow and Keras. It is available at the Github repository: https://forgemia.inra.fr/raphael.mourad/deepgnn

## 3 Results and Discussion

### 3.1 Conservation of protein binding sites in related species

The working hypothesis of the proposed semi-supervised model is the following: transcription factor binding sites as encoded by a DNA motif (or multiple motifs) are evolutionary conserved, meaning that the position of a DNA motif in a genome is conserved in evolutionary close species. This hypothesis implies that DNA sequences in unlabeled evolutionary close genomes could be used to improve predictions.

To verify this working hypothesis, we assessed whether the CTCF binding sites (as encoded by the CTCF DNA motif MA00138 from Jaspar database) in the human genome are conserved in evolutionary close genomes. For this purpose, we binned the human genome (hg38 assembly) into 200 b bins. Then, we liftovered the human genome bins to other mammalian genomes to obtain homologous bins as detailed in Subsection Materials and Methods, Homologous DNA sequences. Then, for each bin of the human and the corresponding homologous bins in the other mammalian genomes, we scanned the DNA sequences for the presence of a CTCF motif. We illustrated the results with a genomic loci of the ERCC4 gene in Figure 2. In this loci, three CTCF binding peaks were found in the human genome (hg38). For each peak, a CTCF binding motif was found in the human genome. By looking at other mammalian genomes, we found that, for each human peak, the CTCF binding motif was also present in apes (chimpanzee: panTro6 and gorilla: gorGor6). By looking at more distantly related species, we still could find the DNA motif for some peaks. For instance, for the peak at the left side, the DNA motif was also conserved in the macaque (rheMac10), the mouse (mm10), the pig (susScr11) and the cow (bosTau9), but absent in the dog (canFam3), the cat (felCat9) and the rabbit (oryCun2).

**Figure 2.**
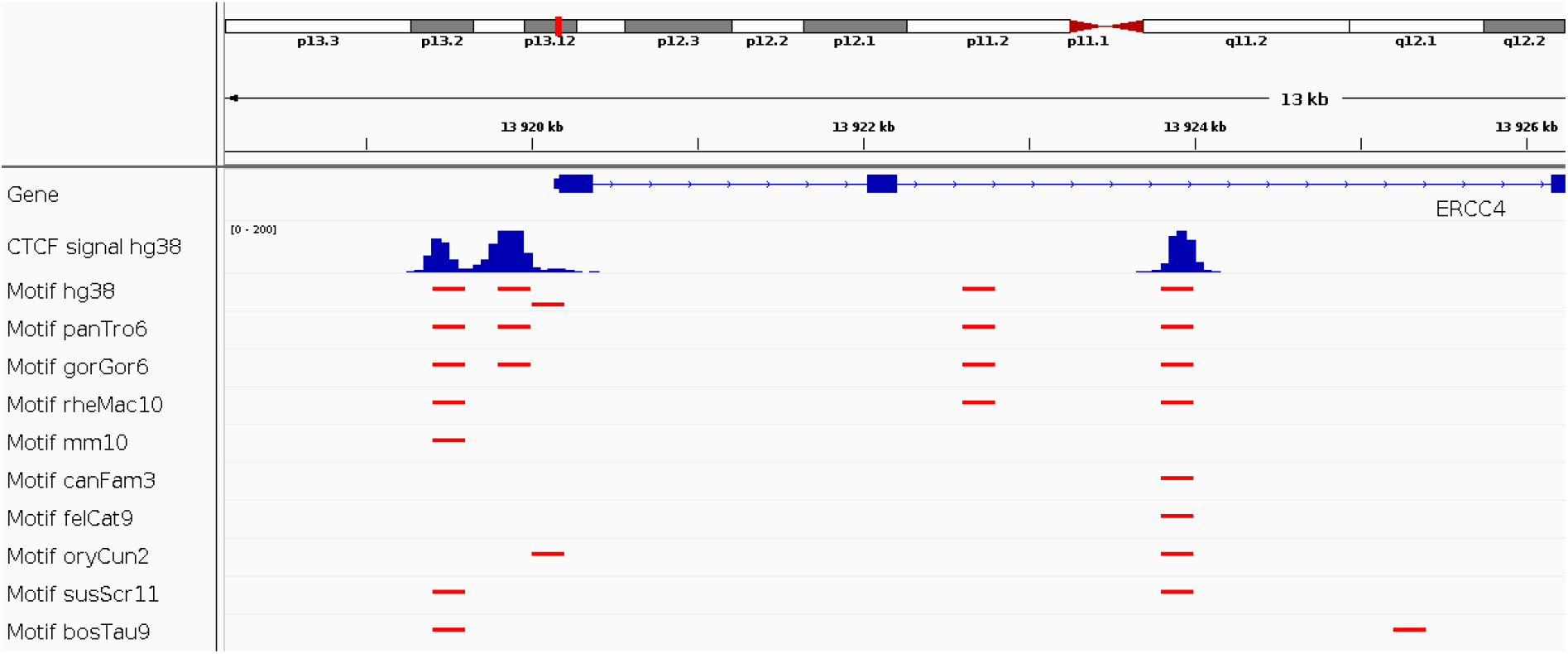
Conservation of CTCF DNA binding sites as predicted from the CTCF motif.

We concluded that using unlabeled genomes could be used to improve the prediction of regulatory sequences by account for the homology of sequences between related genomes since DNA motif position is often conserved.

### 3.2 Prediction of regulatory elements

The proposed model takes as input sets of homologous 1 kb DNA sequences between species and a set of graphs of homologies between the DNA sequences encoded as a binary matrix. The model then predicts the binding of a given TF within a 200 b DNA sequence centered on the 1 kb DNA sequence. The model can predict the binding as a classification problem (whether bin overlaps a binding peak) or as a regression problem (the ChIP-seq coverage signal).

We binned the human genome into bins of 200 b, considered as either bound by a TF (if overlapping > 50%), or not bound otherwise. For each 200 b bin, we extracted the surrounding DNA sequence of 1 kb (containing the bin and the context). We also mapped homologous DNA sequences in other mammal genomes using the liftover program (see more details in Subsection Materials and Methods, Homologous DNA sequences). To train the model, we used bins from chromosomes 1 to 15, and to validate the model, we used bins from chromosomes 16 to 22 and X.

### 3.3 Comparison with the baseline model

We then comprehensively compared the proposed semi-supervised model (CNN-GNN) with the baseline model (CNN) where graph convolution was not used. For a fair comparison between the two models, no fine-tuning was used to favor any of the two models, and the same hyperparameters were used for both models. We first built a semi-supervised model with only one convolutional layer (shallow CNN), with the equivalent baseline model. We trained the models to predict the binding of CTCF from ChIP-seq data as a binary classification setting (peak versus no peak). In term of area under the curve (AUROC), we found a moderate increase with the CNN-GNN when trained with sequences from 23 mammalian genomes (1 labeled genome plus 22 unlabeled genomes) compared to the CNN (from 0.955 to 0.958, +0.3%, Figure 3A). However, genomic data are highly imbalanced with few positive labels (peak presence) compared to negative labels (peak absence), which makes the AUROC a less appropriate metrics to evaluate prediction performance. When we instead compared the area under the precision-recall curve (AUPR), a better metric to account for imbalanced data, we instead observed a strong increase for the CNN-GNN compared to the baseline CNN (from 0.224 to 0.272, +20.8%, corresponding to a raw increase of 0.047, between CNN-GNN with 23 mammals and baseline CNN, Figure 3B). Interestingly, we observed that the AUPR monotonically increased with the number of mammalian genomes used in the CNN-GNN, demonstrating that the higher number of mammalian genomes included, the better prediction. In a regression setting, where the models were trained to predict the CTCF ChIP-seq signal within a bin, we also found a strong increase of Pearson correlation for the CNN-GNN with 23 mammals compared to the baseline CNN (from 0.296 to 0.358, +20.9%, corresponding to a raw increase of 0.062; Figure 3C).

**Figure 3.**
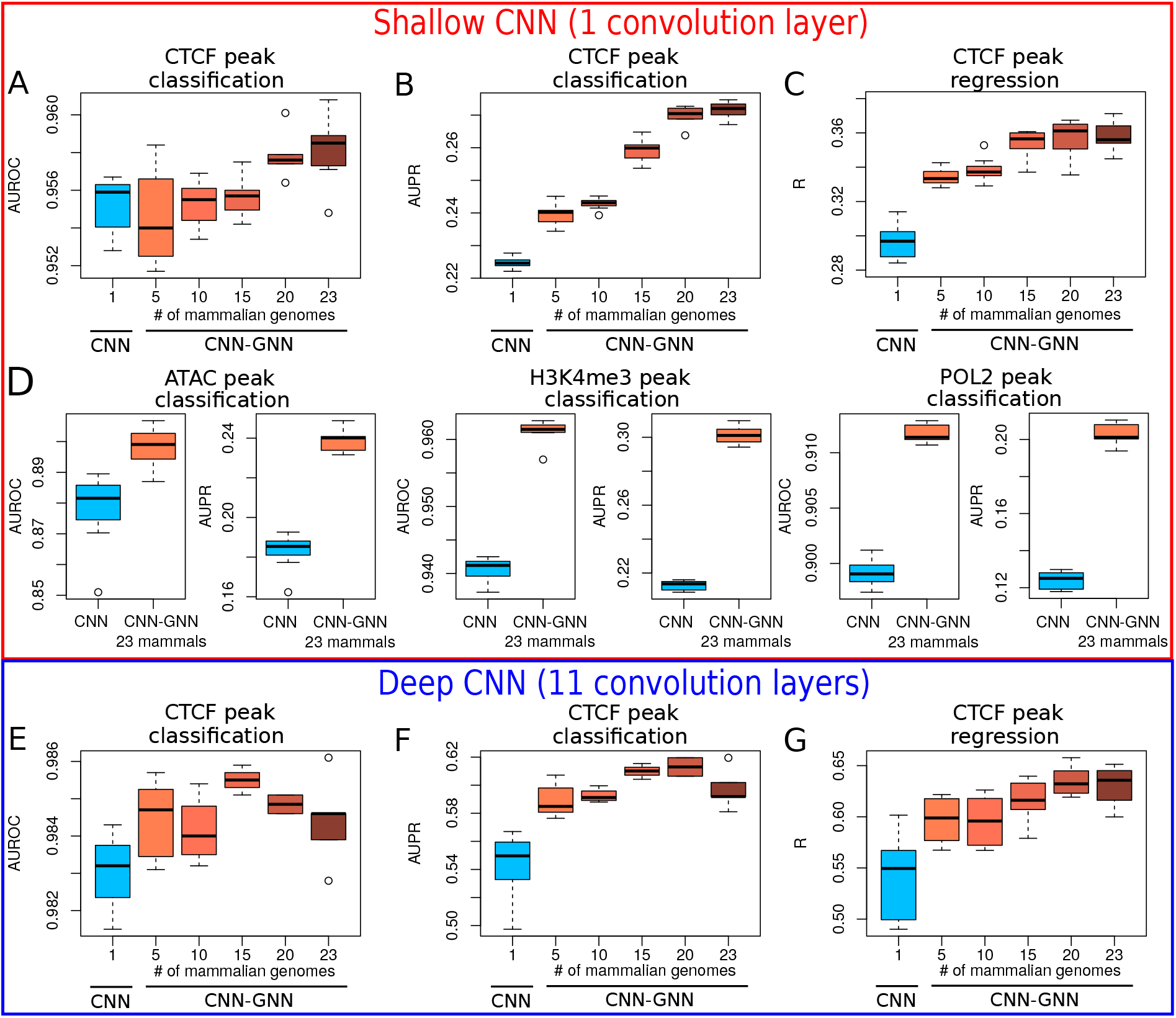
Comparison of prediction performances between the semi-supervised model (here called CNN-GNN) with the baseline model (CNN) where graph convolution was not used. Ten trainings were done for each model to make boxplots. A) Comparison of shallow CNN and shallow CNN-GNN in a classification setting, in term of area under the roc curve (AUROC). B) Comparison of shallow CNN and shallow CNN-GNN in a classification setting, in term of area under the precision recall curve (AUPR). C) Comparison of shallow CNN and shallow CNN-GNN in a regression setting, in term of Pearson correlation. D) Comparison of shallow CNN and shallow CNN-GNN in a classification setting for ATAC peaks, H3K4me3 peaks and POL2 peaks. E) Comparison of AUROC between deep CNN and deep CNN-GNN in a classification setting. F) Comparison of AUPR between deep CNN and deep CNN-GNN in a classification setting. G) Comparison of Pearson correlation between deep CNN and deep CNN-GNN in a regression setting.

We then assessed prediction performance with other types of data, including ATAC-seq, H3K4me3 and POL2 ChIP-seq data (Figure 3D). For ATAC-seq, AUROC was moderately increased (from 0.878 to 0.898, +2.3%), while AUPR was strongly increased (from 0.183 to 0.239, +30.5%). For H3K4me3, AUROC was increased from 0.941 to 0.961 (+2.2%), while a very important increase of AUPR was observed (from 0.213 to 0.301, +41.8%). For POL2 ChIP-seq, AUROC was moderately increased (from 0.899 to 0.912, +1.4%), while AUPR was also moderately increased (from 0.124 to 0.203, +63.5%).

We also sought to assess if our semi-supervised approach could improve the performance for other CNN architectures, for instance when plugged on the top of a deeper CNN architecture. For this purpose, we used a deeper CNN architecture where one convolutional layer was followed by 10 dilated convolutional layers with residual connections similar to [9]. Similarly to the shallow CNN, the CNN-GNN did not improve significantly the classification performance in terms of AUROC (from 0.983 to 0.984, +0.1%; Figure 3E). However, in term of AUPR, the CNN-GNN improved the CNN (from 0.544 to 0.597, +9.9%; Figure 3F). In the regression setting, the CNN-GNN greatly improved the CNN (from 0.541 to 0.631, +16.6%; Figure 3G).

Different graph neural network convolutions were compared with our model implementing a Graph-SAGE convolution (Supp Fig S1). In term of AUROC, we found that the best performances were obtained on average with GraphSAGE (0.958), compared to those obtained with GCN (0.953), GAT (0.955), APPNP (0.957) and GSC (0.957). In term of AUPR, we also found the best performances on average with GraphSAGE (0.272), compared to GCN (0.259), GAT (0.257), APPNP (0.261) and GSC (0.261).

We thus concluded that our semi-supervised approach borrowing information from unlabeled genomes could greatly improves the performance of predictions in many different situations, including classification and regression, shallow and deep CNNs, and transcription factor binding (CTCF), chromatin accessibility (ATAC), histone mark (H3K4me3) and polymerase II binding (POL2).

### 3.4 Cross-species prediction

Previous studies have shown that the grammar of regulatory sequences can be transferred across species, for instance between human and mouse, to make accurate tissue-specific predictions [21]. In fact, crossspecies prediction could be used to annotate the genome of many mammalian species for which no functional data are available, but for which a large body of human data is available. Hence, we explored if our CNN-GNN could improve predictions across species, compared to the baseline CNN. For this purpose, we first trained our model on human CTCF ChIP-seq data and predicted on the mouse genome. In a classification setting, we found that a shallow CNN-GNN trained from human labeled genome and 9 unlabeled genomes slightly improved AUROC (from 0.965 to 0.967, +0.3%), but strongly increased AUPR (from 0.169 to 0.211, +24.7%) (Figure 4A). Similarly, a deep CNN-GNN slightly improved AUROC (from 0.991 to 0.992, +0.1%), but significantly increased AUPR (from 0.508 to 0.588, +15.8%) (Figure 4B). We also explored other functional data. For H3K4me3 classification, we found that a shallow CNN-GNN improved AUROC (from 0.893 to 0.918, +2.8%), but significantly increased AUPR (from 0.274 to 0.321, +17.1%) (Figure 4C). For POL2 classification, we found that a shallow CNN-GNN slightly improved both AUROC (from 0.866 to 0.878, +1.3%) and AUPR (from 0.218 to 0.242, +10.1%) (Figure 4D).

**Figure 4.**
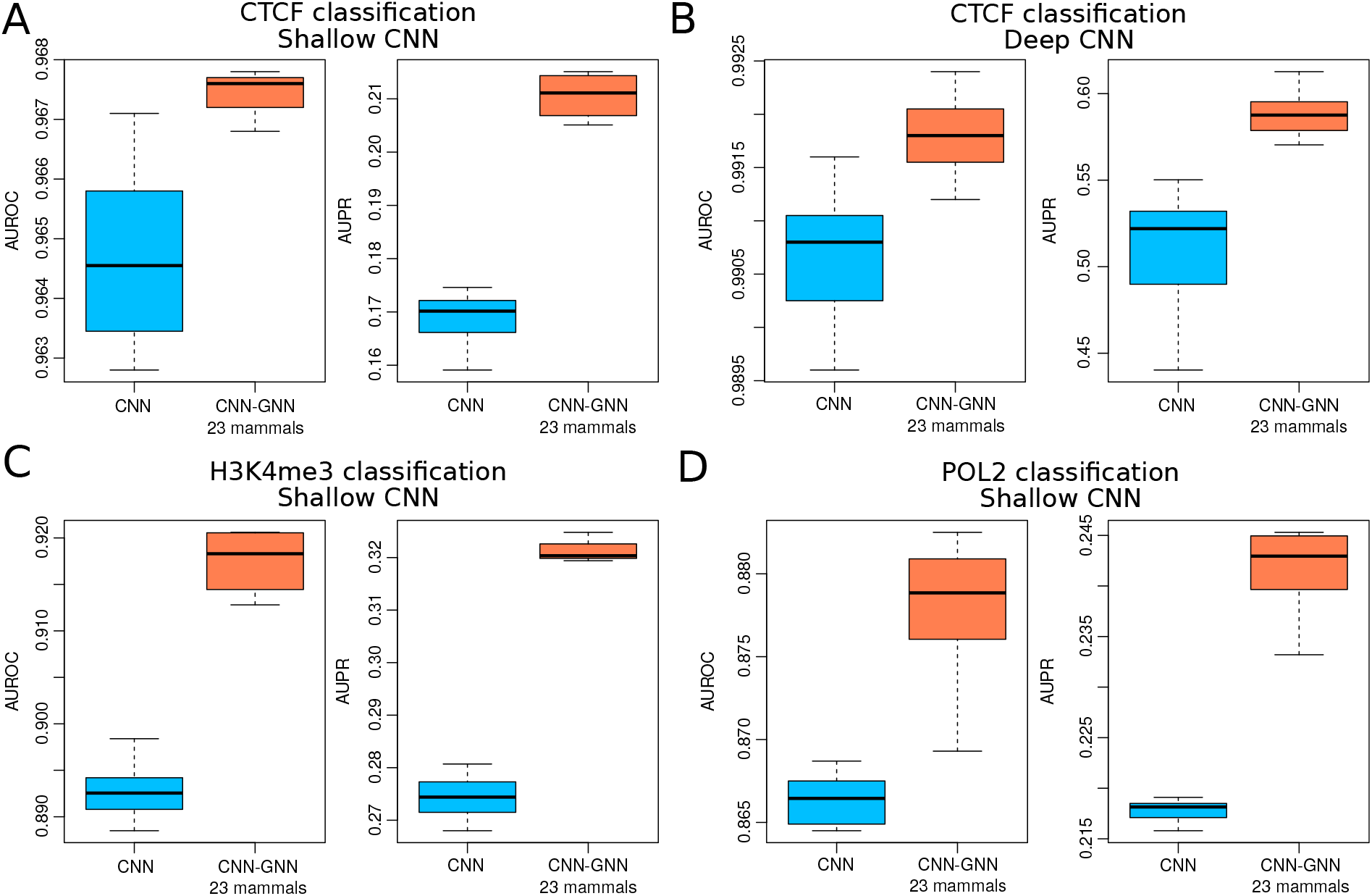
Comparison of prediction performance of our semi-supervised model trained from the human labeled genome and 9 unlabeled genomes to predict mouse experiments, as compared to the baseline model. A) CTCF classification with shallow CNN. Area under the ROC curve (AUROC) and area under the PR curve (AUPR) are plotted. B) CTCF classification with deep CNN. C) H3K4me3 classification with shallow CNN. D) POL2 classification with shallow CNN.

Results thus showed the superiority of our semi-supervised approach as compared to supervised learning for cross-species predictions in mammals for both shallow and deep CNNs, and for transcription factor binding (CTCF), histone mark (H3K4me3) and polymerase II binding (POL2).

### 3.5 Multigenome training compared to semi-supervised learning

Multi-genome training was previously proposed to improve the training of deep learning models [21]. Multi-genome training consists in training a model not only from one labeled genome (for instance from human genome), but from multiple labeled genomes (for instance from the human and the mouse genomes at the same time) followed by fine-tuning on the target labeled genome. Multi-genome training is a simple approach to improve predictive performance, but it is limited by the availability of functional data from multiple genomes. For mammals, most of the functional data are available in human, and to a certain extent in mouse. In comparison, our semi-supervised learning instead allows to exploit many more genomes, *e.g.* all available mammalian genomes sequenced.

We compared multi-genome training (MGT) with our semi-supervised learning (SSL) and one-genome training (OGT) for both shallow and deep CNNs (Figure 5). For shallow CNNs, we observed a slight improvement of MGT compared to the OGT in term of AUROC (from 0.955 to 0.957, +0.2%; Figure 5A), also for SSL with 23 mammals compared to the OGT (from 0.955 to 0.958, +0.3%). However, in term of AUPR, we found a slight improvement for MGT (from 0.224 to 0.230, +2.5%; Figure 5B), but a strong increase for SSL with 23 mammalian genomes as compared to OGT (from 0.224 to 0.272, +20.8%). For deep CNNs, we observed an improvement of MGT compared to the OGT in term of AUROC (0.983 to 0.985, +2.4%; Figure 5C), and almost no improvement of SSLwith 23 mammalian genomes compared to OGT (from 0.983 to 0.984, +0.1%). In term of AUPR, we found an increase for MGT compared to OGT (from 0.543 to 0.572, +5.3%; Figure 5D), and a stronger increase for SSL compared to OGT (from 0.544 to 0.597, +9.9%).

**Figure 5.**
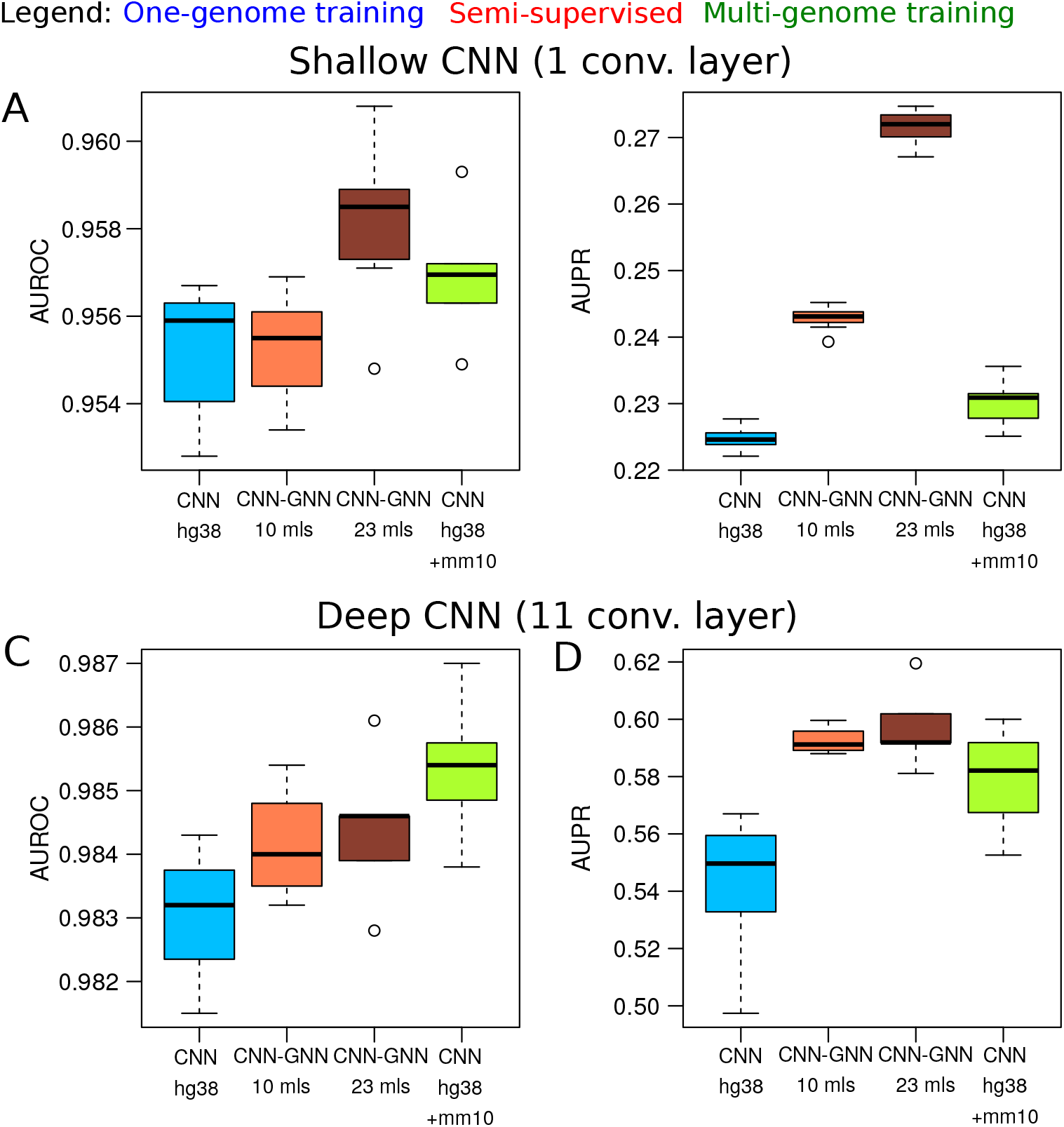
Comparison of prediction performance of our semi-supervised model trained from labeled human genome and 22 unlabeled mammalian genomes with the baseline model trained from human and mouse labeled genomes (also called multi-genome training [21]), and the baseline model trained only from human (one-genome training). A) Area under the ROC curve (AUROC) from shallow CNNs. The baseline CNN trained on labeled human data (hg38), the baseline CNN trained on both labeled human and mouse data (hg38+mm10), and the CNN-GNN trained on 5 or 23 mammalian genomes were compared. B) Area under the PR curve (AUPR) from shallow CNNs. C) Area under the ROC curve (AUROC) from deep CNNs. D) Area under the PR curve (AUPR) from deep CNNs.

These results thus revealed that semi-supervised learning with more than 20 unlabeled genomes provides similar or better performances than multi-genome training with two labeled genomes, while the former does not necessitate any additional functional data as compared to the latter.

### 3.6 Prediction of SNP effects

An important application of deep learning models in genomics is to predict the impact of a SNP on functional data. For instance, studies have shown that deep learning can predict the impact of a SNP on gene expression, histone modification, or protein binding [6, 7, 10]. Hence, we explored the ability of our CNN-GNN to improve the prediction of SNP effect on functional data, and compared it to the baseline CNN. For this purpose, we first trained a deep CNN for CTCF peak classification, and then predicted the impact of a SNP on CTCF binding, as done in [7]. We found a good Pearson correlation between the observed effect as estimated by ChIP-seq allelic imbalance (from ADASTRA database [22]), and the predicted effect (R= 0.312; Figure 6A). We then trained a deep CNN-GNN and observed an increase of Pearson correlation (R= 0.458, +46.8%; Figure 6B). Repeating the same experiment with different model training runs led to the same conclusion (p=9 × 10^-11^; Figure 6C). We also trained a deep CNN for ESR1 peak classification, and then predicted the impact of a SNP on ESR1 binding. We found a moderate Pearson correlation between the observed effect and the predicted effect (R= 0.150; Figure 6D). With the CNN-GNN, we observed an increase of Pearson correlation (R= 0.256, +70.7%; Figure 6E). Repeating the same experiment also showed a significant difference (p=0.03; Figure 6F).

**Figure 6.**
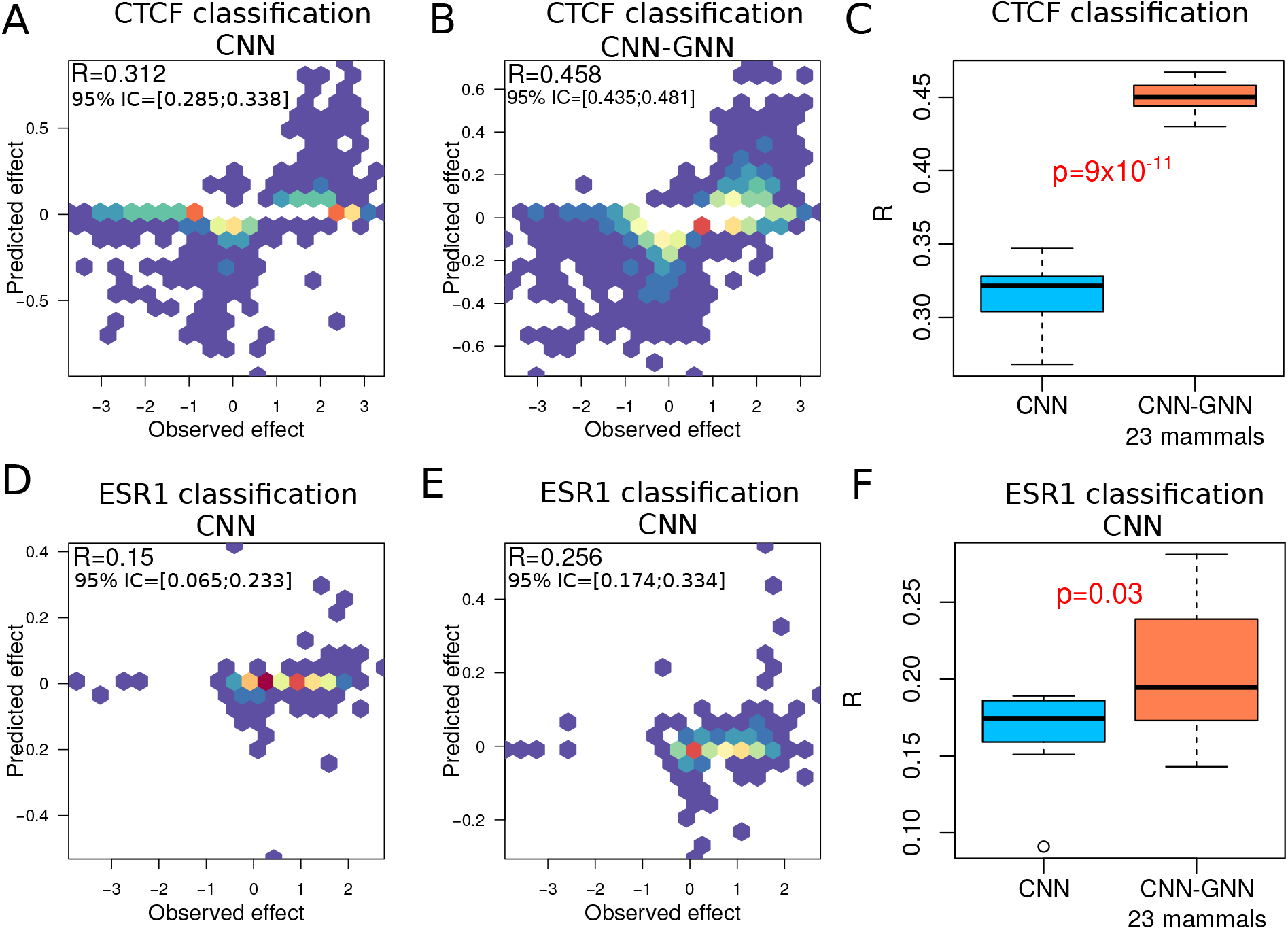
CTCF SNP effect prediction with our semi-supervised model trained from human labeled genome and 22 unlabeled genomes, compared to the baseline model. Allelic imbalance CTCF SNP were downloaded from ADASTRA database. A) Scatter plot of predicted effect versus observed effect with a shallow CNN. B) Scatter plot of predicted effect versus observed effect with a shallow CNN-GNN.

We thus found that our semi-supervised learning could improve the prediction of SNP effect on functional data, and could thus be used to prioritize SNPs in clinical studies.

## 4 Conclusion

In this article, we propose a novel framework for regulatory sequence prediction using semi-supervised learning with graph neural network. For this purpose, neural network training does not only exploit labeled sequences (*e.g.* human genome with ChIP-seq experiment), but also unlabeled sequences available in much larger amounts (*e.g.* from other species without ChIP-seq experiment, such as chimpanzee). For this purpose, a graph neural network is built on the top of a classical convolutional neural network in order to propagate feature information from homologous unlabeled sequences. The framework is flexible and can be plugged into any neural architecture including shallow and deep networks, for which the framework shows predictive performance improvements in both classification and regression tasks. Moreover, improvements were found not only for CTCF ChIP-seq whose DNA motif is known to be conserved among mammals, but surprisingly also for ATAC-seq, H3K4me3 ChIP-seq and POL2 ChIP-seq, which clearly demonstrates the versatility of our approach.

A strong drawback of supervised learning for DNA sequence prediction in human and most species is the limit of the genome size, *e.g.* 3.3 Gb in human. Of course, accounting for variants can increase the amount of DNA sequences, but still represent only a small amount of DNA sequence variation. Multigenome training was previously proposed to exploit labeled sequences from multiple genomes during model training, allowing in a simple manner to train from a larger DNA sequence space [21]. While such approach shows good performance improvements, it requires the availability of many labeled genomes for a given functional experiment, which are most of the time not available. Semi-supervised learning further extend this philosophy by exploiting during training DNA sequences from genomes that are not even labeled, which represents the majority of genomes. Semi-supervised learning was successfully applied for cross-species prediction, allowing the annotation of genomes in species without functional data. Moreover, we found that such approach could improve the prediction of the impact of SNPs and thus represent an interesting avenue for human genetic research.

There are several limitations of the proposed approach. First, the use of our approach is likely limited to species which are not too evolutionary distant. For instance, it is very unlikely that including plant genomes would help to predict regulatory sequences from a mammalian genome. Second, we did not try to plug our GNN on the top of other networks such as the transformer network, and thus do not know whether our approach could bring improvement. Third, our approach provides improvement without any additional functional data in homologous species, but as a drawback it requires higher computational resources.

## Authors’ contributions

RM conceived and designed the project. RM implemented the model and analyzed the data. RM wrote the manuscript.

## Acknowledgements

The author is grateful to Nathalie Vialaneix and Céline Brouard, and the SaAB team from INRAE MIAT lab.

## Supplementary Figures

**Supp Fig S1:**
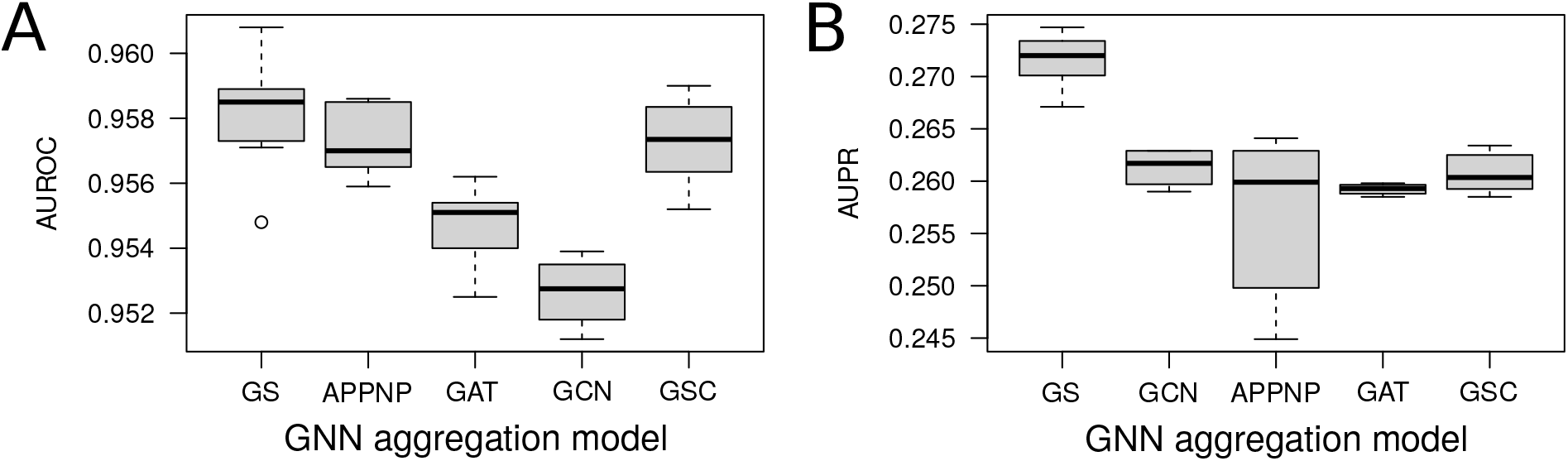
Comparison of prediction performances between different graph neural network convolution layers. A) Comparison in term of area under the roc curve (AUROC). B) Comparison in term of area under the precision recall curve (AUPR).

